# Host response and cell fates of human U937 monocytes upon infection with vancomycin-resistant *E. faecium*

**DOI:** 10.64898/2026.10.20.760358

**Authors:** Annika Walter, Theresia Prenzel, Thorsten Bischler, Bärbel Kieninger, Iana Gadjalova, Tom Gräfenhan, Wulf Schneider-Brachert, Anca Rath, Jürgen Fritsch

## Abstract

In recent years, *Enterococcus faecium* has become a high-priority human opportunistic pathogen, especially due to its antibiotic and environmental resistance, promoting hospital-acquired infections. However, its interaction with host cells remains poorly understood.

In this study, we investigated the interaction of four recently collected, clinical vancomycin-resistant *Enterococcus faecium* (*VREfm*) isolates with the human U937 cell line. Early after infection, we observed close bacterial association with U937 monocytes and features suggestive of internalization. We thus investigated host cell fates after washing at 2 hours post infection (hpi) to concentrate on effects induced by internalized bacteria or those with direct host cell contact.

Whereas viability was not impaired at 6 hpi, moderate cell death was induced to an isolate-specific extent at 24 hpi, which was abrogated when direct host-pathogen contact as prevented via boyden chamber cell culture inserts, but was enhanced by TNF-R1 deficiency.

Host mRNA sequencing revealed a consistent pattern of differentially expressed genes across all *VREfm* isolates, suggesting a shared innate immune and stress-related response at 6 hpi, with functionally enriched biological processes related to leukocyte activation, cytokine signaling, programmed cell death, and RNA stability.

Together, these data show that *VREfm* triggers an early pro-inflammatory transcriptional response in U937 monocytes without immediate cytotoxicity in U937 monocytes, while late-stage isolate-specific cell death induction depends on direct host cell contact or internalization and involves TNF-R1 mediated regulation. Our findings may assist future efforts to understand how *VREfm* triggers host cell damage that contributes to severe inflammation and systemic infection such as persistent bacteremia.

**One-Sentence Summary:** Vancomycin-resistant *Enterococcus faecium* closely associates with U937 monocyte membranes and displays features suggestive of internalization, triggering early proinflammatory transcriptional responses and later, isolate-specific cytotoxicity that requires direct host-pathogen contact and is modulated by TNF-R1.

## Introduction

*Enterococcus faecium* (*Efm*) is a Gram-positive opportunistic pathogen that is not only part of the human intestinal microbiome, but has emerged as a major cause of hospital-acquired infections worldwide. Its remarkable ability to acquire antibiotic resistance, particularly to vancomycin (vancomycin-resistant *E. faecium*: *VREfm*), has led to its designation as a high-priority pathogen on the World Health Organization (WHO) list for research and development of new antibiotics (WHO, 2024). Upon translocation across protective tissue barriers, severe systemic infection can occur, particularly life-threatening bloodstream infections (BSIs), whose incidence has increased in recent years, especially among elderly and immunocompromised patients (Ayobami et al., 2020; Brinkwirth et al., 2025; Canadian Nosocomial Infection Surveillance, 2026; Leding et al., 2026). Corresponding epidemiology and underlying virulence mechanisms are becoming increasingly well studied, whereas comparatively little is known about the molecular basis of host-pathogen interactions during *Efm* infection, impeding the progress toward targeted therapeutic or prevention strategies. Addressing this gap requires a deeper exploration of *Efm* virulence factors and their role in modulating host cellular responses (Wei et al., 2024).

Several human pathogens, including members of the ESKAPE group (e.g., *S. aureus, K. pneumoniae, A. baumannii*, or *P. aeruginosa*) exploit a facultative intracellular lifestyle to create a favorable, protective niche in hostile host environments (Cai et al., 2025; Szperlinski et al., 2025; Thakur et al., 2019). This strategy can promote disseminating infection through uptake by intestinal barrier cells or professional phagocytes, followed by intracellular proliferation and eventual bacterial release (Wei et al., 2024), and has also been described for the related pathogen *Enterococcus faecalis* (*Efc*) (Archambaud et al., 2024). Although immune evasion and adherence to host cells and foreign bodies are well-recognized mechanisms by which *Enterococcus faecium* (*Efm*) adapts to the host environment (Cairns et al., 2023), an intracellular lifestyle has rarely been associated with this pathogen.

Underlying molecular mechanisms and specific proteins involved in the pathogen-host dialogues may however represent novel targets for the prevention and treatment of infectious diseases, particularly in case of common pathogens formerly described as merely extracellular.

We previously investigated the spread of vancomycin-resistant enterococci in clinical settings, revealing ST117, ST80, and ST1299 as the most prominent patient-associated strains in southeast Germany (Caplunik-Pratsch et al., 2024; Rath et al., 2025; Rath and Schneider-Brachert, 2026). In the present study, we investigated cross-talk between four selected *VREfm* isolates and the human U937 monocyte cell line, showing binding to and internalization by host cells across isolates, contact-dependent and TNF-R1-regulated cytotoxicity, along with a shared core of differentially expressed host transcriptional pathways related to inflammatory cell activation, proliferation, and survival.

## Methods

### Human cell lines, bacterial strains, and culture conditions

U937 cells were obtained from DMSZ and cultivated in RPMI (Gibco) supplemented with 10% FBS (Gibco) and 1% sodium-pyruvate (Gibco) without antibiotics at 37°C, 5% CO_2_. The generation of U937Δ_TNF-R1_ cells was described before (Zingler et al., 2019). Vancomycin-resistant *E. faecium* (*VREfm*) strains VRE 23×1609 (V2: SAMN61876551), VRE 23×1608 (V3: SAMN61876552), VRE 7282 (V6: SAMN61876553), and VRE 8170 (V7: SAMN61876554) (collected in 2022 or 2023; names according to in-house culture collection; Table 1) were grown in brain-heart infusion (BHI) medium aerobically at 37°C and 180 rpm overnight. The bacterial overnight cultures were adjusted to the stationary growth phase at an optical density at 600 nm (OD_600_) of 1.0–1.5 following incubation for at least 1h at 37°C and 180 rpm. To calculate the multiplicity of infection (MOI), the bacterial number was determined with an OD_600_ of 1 corresponding to 8 × 10^8^ colony-forming units (CFU) per ml. By default, VRE were grown on blood agar to observe colony morphology and growth differences.

**Table 1.** Overview of vancomycin-resistant *Enterococcus faecium* isolates used in the present study.

| Isolate name | 23x1609 | 23x1608 | 7282 | 8170 |
| --- | --- | --- | --- | --- |
| Alternative name | V2 | V3 | V6 | V7 |
| Sequence Type | ST80 | ST117 | ST1299 | ST1299 |
| BioSample accession number | SAMN61876551 | SAMN61876552 | SAMN61876553 | SAMN61876554 |

### Genomic sequencing of VREfm isolates

We performed Illumina short-read sequencing as previously described (Rath et al., 2025). We performed long-read sequencing using Oxford Nanopore Technologies with the Ligation Sequencing Kit V14. Hybrid genome assemblies were generated using MiLongA v1.0.3 (Deneke, 2020) with default parameters, and the Flye v2.9.4-b1799 assemblies produced within the pipeline were used for all downstream analyses. Genome annotation was performed using the NCBI Prokaryotic Genome Annotation Pipeline (PGAP; release 2023-10-03.build7061) (Tatusova et al., 2016). Gene set similarity was quantified using the Jaccard coefficient (intersection/union of genes) in R (v4.5.1; RStudio v2025.5.1.513) and visualized as a heat map using the *pheatmap* package. Presence or absence of genes per isolate is listed in **Supplementary File S1**.

### RNA sequencing and data analysis

To screen for differential gene expression upon infection, U937 cells were infected with four *VREfm* isolates for a total of 6 h, including a washing step at 2 hours post infection (hpi), or left untreated for 4 h. RNA isolation, library preparation, sequencing and data analysis were performed together with *S. aureus*-infected cells as detailed before (Walter et al., 2026). Library preparation and mRNA-seq were performed at the Genomics Core Facility “KFB - Center of Excellence for Fluorescent Bioanalytics” (University of Regensburg, Regensburg, Germany www.kfb-regensburg.de). Processing of raw data was performed at the Core Unit Systems Medicine, University of Würz burg (Wurzburg, Germany; https://www.med.uni-wuerzburg.de/cu/sysmed/).

For downstream visualization and functional analyses, gene lists were defined as protein-coding, significantly regulated (adjusted p-value after Benjamini-Hochberg correction (padj) < 0.05), and at least two-fold up- or downregulated (| log2FC ≥) relative to uninfected control cells. Full STRING networks (https://string-db.org/) were generated at medium confidence (0.400), with edges indicating confidence and disconnected nodes removed.

Principal component analysis (PCA) was performed by applying variance-stabilizing transformation (vst) to normalized read counts via DESeq2. The 1000 most variable, protein-coding genes (after removal of zero/NA-variance features) across all samples were selected and scaled prior to PCA computation using R *(preamp*, stats package). Samples were grouped by condition (VRE 23×1608, VRE 23×1609, VRE 7282, VRE 8170, and untreated) and PCA scores were plotted using ggplot2 (version 3.5.2). Z-score heat maps with gene-wise z-scaling were generated using the *pheatmap* package in R with hierarchical clustering (rows: correlation distance; columns: Euclidean distance; complete linkage). R version 4.5.1, RStudio 2025.5.1.513.

Functional enrichment of biological pathways present in these gene lists was analyzed using Metascape version *v3*.*5*.*20260701* (Zhou et al., 2019). We used a list of 19900 protein-coding genes from RNA sequencing of VREfm- and S. aureus-infected U937 cells as the enrichment background. GO Biological Processes (GO:BP) was used as the ontology source for multiple gene lists, and terms with a p-value < 0.01, a minimum count of 3, and an enrichment factor > 1.5 were included and grouped into clusters based on their membership similarities. The overlaps between the isolate-specific gene lists and between genes sharing the same enriched ontology term(s) are visualized by Circos plots via Metascape.

The cluster is represented by the driving member with the lowest padj-value. Log1O(P) values, the number of involved genes and pathways per cluster were extracted and visualized as bubble plots in R using *ggplot2 (version 3*.*5*.*2)*. Full DESeq2 lists and enrichment results are available in **Supplementary File S1**.

### Infection assays

U937 cells were adjusted to l×l0^6^ cells/ml prior to infection and infected with live *VREfm* at MOI 50, followed by 2 h of co-culture at 37°C, 5% CO_2_ to allow for synchronized intracellular uptake. To remove planktonic extracellular bacteria, we washed cells and tissue culture wells with PBS (G1397; Sigma-Aldrich) to concentrate, though not limit, the investigation of infection-induced effects by surface-bound or intracellular *VREfm*. For imaging approaches (Imaging flow cytometry and Giemsa staining), gentamicin was added at 500µg/ml after the washing step to help eliminate remaining extracellular bacteria, although gentamicin susceptibility varied between isolates.

For cell-death experiments, bacteria were either added directly to the cells and mixed by gentle shaking or applied to Boyden chamber cell-culture inserts (0.4 µm 83.3932.041, Sarstedt) placed into the wells of the tissue-culture plate to prevent direct host-pathogen contact.

*Annexin V/7-AAD staining:* at the indicated time points, 250 µl of carefully resuspended U937 cells were sedimented (500 x g, 5 min), washed twice, resuspended in 20 µl staining mix (18 µl PBS, 1 µl FITC-Annexin V, 1 µl 7-amino-actinomycin D (7-AAD); #640922, Biolegend), and incubated for 20 min in the dark. 100 µl of binding buffer were added to the samples prior to measurement. Data were acquired using a Guava easyCyte (Cytek) and guavaSoft (4.5.25, Cytek) for data analysis. Cells that stained positive for Annexin V or 7-AAD alone or combined were included in the population referred to as undergoing cell death.

*Western blot:* at the indicated time points, cells were sedimented and washed with PBS (5 min, 500 × g, 4°C). The cell pellets were lysed in 30 µl modified RIPA buffer (50 mM TRIS-HCl [pH 7.5], 150 mM NaCl, 1% NP-40, 1% Triton X-100, 1 mM EDTA, 0.25% Na-deoxycholate), containing a protease inhibitor cocktail (4693124001, Sigma-Aldrich), followed by 20 sec sonication (cooled to 4°C) and 5 min sedimentation at 1500 × g at 4°C. The supernatant was used for BCA protein quantification (Pierce, #23225). Samples (20 µg) were prepared immediately before SOS-PAGE using 12.5% or 10% PAA gels. Proteins were blotted onto a PVDF membrane (200T.1, Carl-Roth). The membranes were blocked with 5% skimmed milk in TBS-T and incubated overnight with the primary antibodies, diluted 1:1,000 in 5% skimmed milk. The peroxidase-conjugated secondary antibodies were incubated for 1 h, diluted 1:10,000 in TBS-T. Blots were imaged using ECL (RPN2232, Cytiva) and the ImageQuant800 device (Cytiva). Primary antibodies (1:1,000): PARP-1 (cs #9542S), cleaved caspase-3 (#9661S, Cell Signaling), caspase-3 (#9668S, Cell Signaling), Actin-HRP (HRP-60008, Proteintech). Secondary antibody: anti-rabbit-HRP at 1:10,000 (111-035-144, Jackson ImmunoResearch).

### Imaging flow cytometry

Bacteria were labeled with 5 µM CFSE/PBS for 1 h, shaking at 37°C, followed by three washing steps (10,000 × g, 10 min) in PBS. l×l0^6^ U937 cells were infected with CFSE-labeled bacteria for 3 h at MOI 50. At 2 hpi, a washing step was performed, followed by the addition of 500 µg/ml gentamicin to help eradicate extracellular bacteria when susceptible. 5 min before the end of the infection period, Cell Mask® deep red membrane stain (C10046; Thermo Fisher) was added, diluted 1:25,000 in PBS, and incubated for 5 min, followed by washing twice in cold PBS. Cells were then fixed in 4% PFA/PBS for 15 min on ice, followed by washing in 1 ml cold PBS. Cells were stored in the dark and cooled until imaging using ImageStream MKII (Cytek). 10,000 images per condition were acquired at 60x using the Inspire software, capturing channels Ch01 (Brightfield), Ch02 (CFSE), and Chll (CellMask deep red). Images were analyzed using Ideas software (V6.2.65): Cells were gated: in-focus/single cells/double positive. Then, composites of Ch02/Chll were generated.

### Giemsa staining

Infected U937 cells were analyzed at 3 hpi, and were washed and treated with gentamicin at 500 µg/ml at 2 hpi. After centrifugation at 500 × g for 5 min, the pellet was resuspended in 50 µl PBS and smeared on a microscope slide, followed by MeOH fixation (5 min, RT), air drying and staining for 15 min using Giemsa stain (1:20 in H_2_O; GS500-500ml, Sigma-Aldrich), washing in H_2_O and air drying (GS500-500ml, Sigma-Aldrich).

### Statistical analysis

Statistical analysis was performed using GraphPad Prism 10.6.1 software (GraphPad Software Inc.). Normality and Long-normality Tests were performed using the Shapiro Wilk test. Two-Way ANOVA with Dunnet’s correction were performed for testing of multiple groups. Comparisons were based on infected vs. uninfected control cells or between infected cells with different isolates, where indicated. A p value of< 0.05 was considered statistically significant.

## Results and Discussion

### Characterization of *VREfm* isolates

Four recent clinical *VREfm* isolates were selected based on previous studies that have revealed ST1299 as the major clinical strain besides ST80 and ST117 in southeastern Germany (Caplunik-Pratsch et al., 2025; Rath et al., 2025): 23 × 1608 (ST117), 23×1609 (ST80), 7282 (ST1299), 8170 (ST1299). Jaccard similarity between the isolate’s genomes show genomic variability across isolates, with clustering of 23×1608 and 13×1609 as well as 7282 and 8170 (**Fig. 1A**). Phenotypically, the isolates revealed similar growth on blood agar plates, except for 23×1609, which exhibited colonies of a distinctly small morphology (**Fig. 1B**) as well as slower growth in broth cultures compared to the other isolates particularly during the first two hours following dilution (**Fig. 1C**)

**Figure 1:**
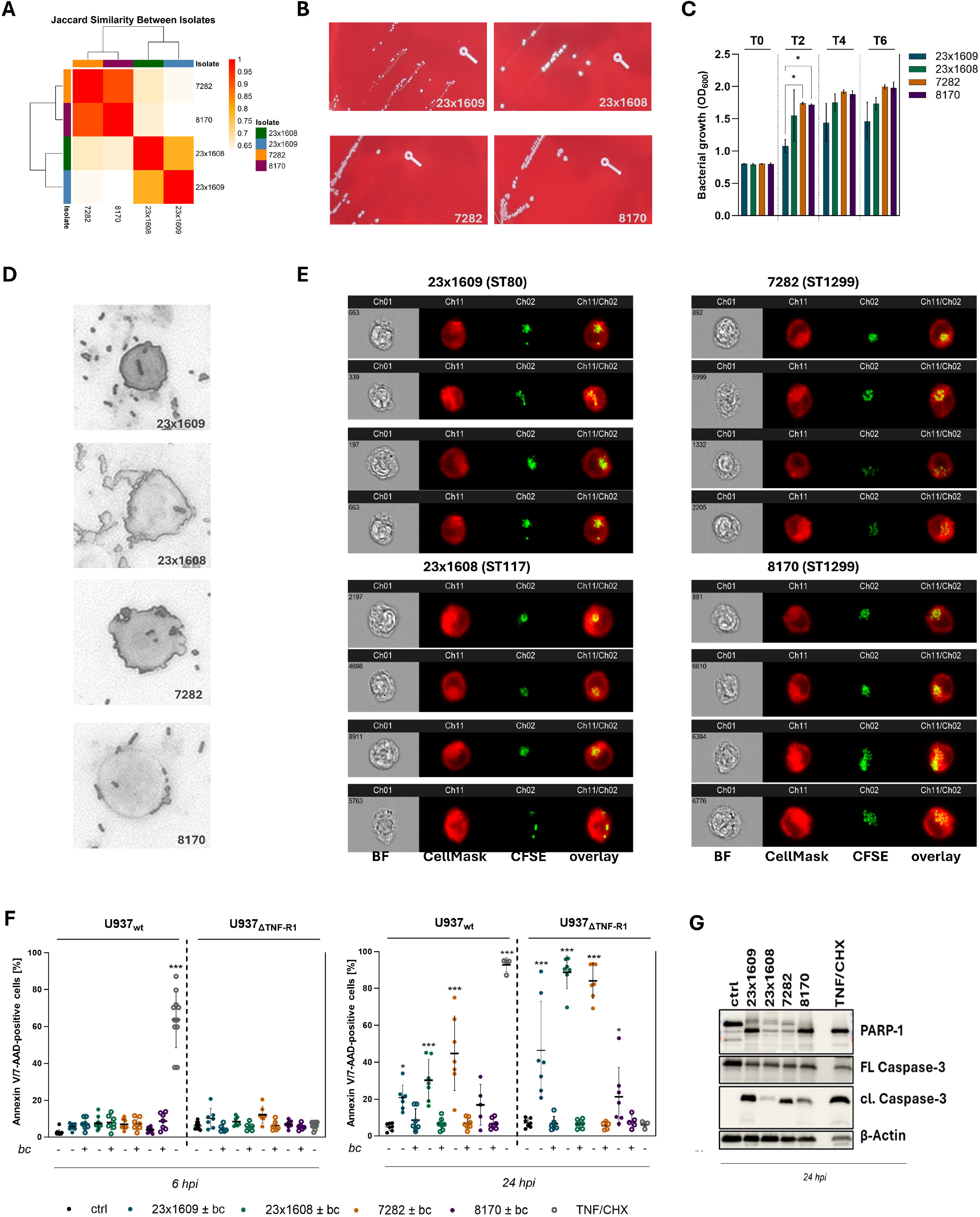
Characterization of *VREfm* isolates and interaction with U937 monocytes at MOI 50. **(A)** Jaccard similarity indices between the genomes of *VREfm* 23×1609, 23×1608, 7282, and 8170, and **(B)** colony morphologies on blood agar showing reduced growth of 23×1609, confirmed by (C) bacterial growth measurements in shaking suspensions during the first 6 hours following dilution. (D) Giemsa staining of infected 11937 cells at 2 hpi shows extracellular, membrane-bound and putative intracellular *VREfm*. **(E)** Close association of *VREfm* with U937 cells and colocalization that could be consistent with internalization was underscored by imaging flow cytometry of infected 11937 cells at 3 hpi. (F) Annexin V/7-AAD staining of infected wild-type and TNF-Rl-deficient U937 cells at 6 hpi (left) and 24 hpi (right) analyzed by flow cytometry shows isolate-dependent cell death induction. **(G)** Activation of apoptotic markers was confirmed by Western blot at 24 hpi. Actin serves as a loading control; representative blots are shown. **C-E** included gentamicin at 500 pg/ml after the washing step at 2 hpi. Quantitative data show mean ± SD of n>3 independent experiments. Statistical analysis was performed using Two-way ANOVA with Dunnet’s correction for Multiple Comparisons. ^*^p<0.05; ^**^p<0.01; ^***^p<0.001 vs. uninfected control, or between isolates if indicated.

This observation indicates a phenotype distinct from that of the other isolates, with features resembling those of small-colony variants (SCVs). Generally, SCVs are highly adapted to a long-term intracellular survival by e.g. a profoundly conversed metabolism, restricted growth, and reduced expression of virulence factors (Kahl et al., 2016). Best described for *S. aureus*, only three reports have identified *Efm* SCVs in both immunocompetent and immunocompromised patients with bacteremia and endocarditis (Egido et al., 2016), sepsis during chemotherapy (Gröbner et al., 2012), and spondylodiscitis (Höring et al., 2019), respectively. For the pathogen’s close relative *Efc*, mounting evidence suggests a proficient intracellular lifestyle in multiple human host cell types (Baldassarri et al., 2005; da Silva et al., 2022; Horsley et al., 2013; Millan et al., 2013; Ogihara et al., 2016; Sussmuth et al., 2000; Wells et al., 2000), but few case reports or *in vitro* studies describe SCV-like phenotypes (Benes et al., 2013; Ogihara et al., 2016; Thieme et al., 2026; Wellinghausen et al., 2009). Given their close association with antibiotic tolerance and resistance, characterizing such phenotypes in clinically important nosocomial pathogens such as *VREfm* is highly relevant. However, this requires analysis beyond the morphological features observed for isolate 23×1609, including auxotrophy testing for NAD and thymidine, as well as investigation of additional isolates from persistent and recurrent infections.

### *VRE/m* closely associates with U937 monocyte membranes and shows features consistent with internalization

Because the initial recognition of a pathogen by innate immune cells is decisive for shaping all subsequent antimicrobial host responses, the ability to disrupt these early sensing pathways provides invasive bacteria with a substantial advantage. For instance, *Efc* stimulates anti-inflammatory cytokine production (e.g. IL-10) to promote immune tolerance (Di Giacinto et al., 2005; Kao and Kline, 2019), impedes opsonization, but is also able to persist in these host cells, which further promote transition from initial infection sites to the bloodstream (Kao and Kline, 2019; Zou et al., 2014). Corresponding mechanisms for *VREfm*, however, are yet to be deciphered.

Considering the pathogen’s high prevalence in BSIs (Cairns et al., 2023; Caplunik-Pratsch et al., 2025), we aimed to investigate these early host-pathogen interactions under controlled and reproducible conditions employing the U937 monocytic cell line, which offers a robust human model system for dissecting the molecular mechanisms governing innate immune recognition.

Giemsa staining revealed the presence of extracellular planktonic and host membrane-bound *VREfm*, but also bacterial cells that appear to be internalized by U937 cells (**Fig. 1D**). Following this path, we show that colocalization of bacteria and host cell staining suggestive of intracellular localization was observable via imaging flow cytometry across all tested *VREfm* isolates at 3 hpi (**Fig. 1E**). However, the definitive identification of bacterial internalization requires additional experimental validation, including adequate imaging approaches such as transmission electron microscopy and colocalization analysis with intracellular compartments, cell fractionation techniques, or the confirmation of viable intracellular colonies when the eradication of extracellular bacteria across all isolates can be ensured.

### U937 host cell fates are modulated by host-pathogen contact and TNF-Rl signaling

Many pathogens, including those with a facultative intracellular lifestyle, manipulate host cell death to a highly isolate- and host cell-dependent extent (Dai et al., 2020; Fraunholz and Sinha, 2012; Li et al., 2019; Walter et al., 2026; Zou and Shankar, 2014). However, how *VREfm* infection affects host cell fates remains poorly understood.

We recently reported U937 cell fates to be critically regulated by the TNF/TNF-R1 axis upon intracellular *S. aureus* infection. Both viruses and bacteria actively modulate TNF/TNF-R1-signaling for their survival at infection sites. For instance, mice that have been fed heat-killed *Efc* showed reduced levels of several cytokines including TNF in an atopic dermatitis model (Choi et al., 2016). A key mechanism by which TNF-R1 activation may contribute to the eradication of internalized bacteria involves its ubiquitination-dependent receptor endocytosis, endo-lysosomal maturation, and fusion (Fritsch et al., 2014). This process enables the activation of anti-microbial enzymes such as cathepsin D (Fritsch et al., 2016; Fu et al., 2020; Heinrich et al., 2004) and aSMase (Edelmann et al., 2011; Schramm et al., 2008), or by coupling TNF-R1 internalization to ROS generation (Morgan and Liu, 2010; Yazdanpanah et al., 2009). Blocking receptor endocytosis can prevent cell death induction, thus maintaining the pathogen’s survival niche (Dreschers et al., 2017; Schneider-Brachert et al., 2006; Waguia Kontchou et al., 2016).

Although the TNF/TNF-R1 system is an established modulator of immune defense against intracellular pathogens, the exact mode(s) of action, especially in different cell/tissue types and the impact of anti-TNF therapy on infection risk, are unclear (Li et al., 2020). Correspondingly, a recent study reported that patients receiving anti-TNF therapy are at increased risk for BSIs at a pathogen-specific level, particularly by *S. aureus*, followed by *Efm* (Leding et al., 2026). Considering the key role for TNF in granuloma formation and intracellular killing, “classical” intracellular pathogens such as *Mycobacterium tuberculosis* or *Listeria monocytogenes* are primarily argued in the context of infections under TNF-inhibition (Leding et al., 2026), but might be of high concern also in ubiquitous pathogens that are less commonly associated with intracellularity, such as *S. aureus* or *Efm*.

In the light of this, we quantified infection-induced cell death by Annexin V/7-AAD flow cytometry after washing at 2 hpi to focus on effects induced by internalized or surface-bound *VREfm* in U937 cells that either express (U937_wt_) or lack TNF-R1 (U937_ΔTNF-R1_).

At 6 hpi, host cell viability was sustained in both U937_wt_ and U937_ΔTNF-R1_ cells (**Fig. 1F, left**), while the positive control TNF/CHX induced apoptosis only in the former, as expected. At 24 hpi, each isolate induced significant cell death to a varying extent, except 8170 showing only a moderate trend towards cytotoxicity. These effects were markedly enhanced in U937_ΔTNF-R1_ cells across isolates by up to two-fold compared to wild-type cells, emphasizing a regulatory role for the TNF/TNF-R1 axis in the finely tuned balance of host defense and survival. Interestingly, preventing direct host-pathogen contact by bc cell culture inserts abrogated host cell cytotoxicity at 24 hpi (**Fig. 1F, right**), indicating that extracellular toxin secretion alone might be insufficient to induce the observed host cell death. This further supports the hypothesis that *VREfm* might employ host cell contact or uptake to modulate host responses.

Western blot analysis confirmed the engagement of pathways involved in apoptotic signaling, as evident from cleavage of PARP-1 and caspase-3 at 24 hpi (**Fig. 1G**). Of note, band intensities of PARP-1 and cleaved caspase-3 were not consistent (apart from 23×1609) and not entirely congruent with cell death intensities quantified by flow cytometry, suggesting additional levels of regulation. The involvement of additional cell death programs such as pyroptosis or necroptosis should be investigated, including caspase-1 activation and gasdermin D cleavage, or MLKL oligomerization, respectively.

To date, a limited number of lytic or host cell homeostasis-disrupting virulence factors have been reported for *Efm* (Fiore et al., 2019), including the botulinum neurotoxin-like toxin (BoNT-like toxin) (Zhang et al., 2018), or enterococcus pore-forming toxins (Epsx) (Xiong et al., 2022). The latter has been shown to lyse HeLa cells and PBMCs in an MHC-I dependent manner. *Efm*-induced lytic cell death has been reported in various human and murine cell lines for certain strains, including U937, while requiring lysozyme treatment (Horing et al., 2012). To unravel tissue adaptation processes, these reports highlight the need to extend our analyses in future experiments to additional non-professional phagocytic host-cell types, differentiated macrophage models, and, in particular, primary cells, together with further *VREfm* isolates derived from different infection sources.

### Homogenous host transcriptional reprogramming across different *VREfm* isolates

To investigate the host response to infection with *VREfm* that is primarily induced by host cell contact or uptake, we aimed to remove planktonic extracellular bacteria at 2 hpi via washing and host mRNA sequencing was performed at 6 hpi, hence prior to the onset of host cell death. Data were acquired in parallel with *S. aureus*-infected and uninfected control cells (*S. aureus* data not shown (Walter et al., 2026).

PCA and heatmap of sample triplicates showed a homogenous, but isolate-specific clustering of samples, except for some variation of one of the 7282 samples, which was still included in downstream analyses (**Fig. 2A+B**). We observed several unique DEGs per isolate (**Table 2**) alongside a shared core of upregulated (N: 43; **Fig. 2C**) and downregulated genes (N: 80; **Fig. 2D**) across the four isolates (padj ≤ 0.05, |log2FC| ≥ 1). **Fig 2E-G** depicts volcano plots including labels for the top 15 differentially expressed genes per isolate. Row-wise z-scored expression values derived from protein-coding vst-data x DEseq2 DEGs (sample (padj ≤ 0.05, |log2FC| ≥ 1) are provided in **Fig. S2**, a selection of 50 DEGs with the highest variance across the included samples in **Fig. S3**.

**Table 2.**
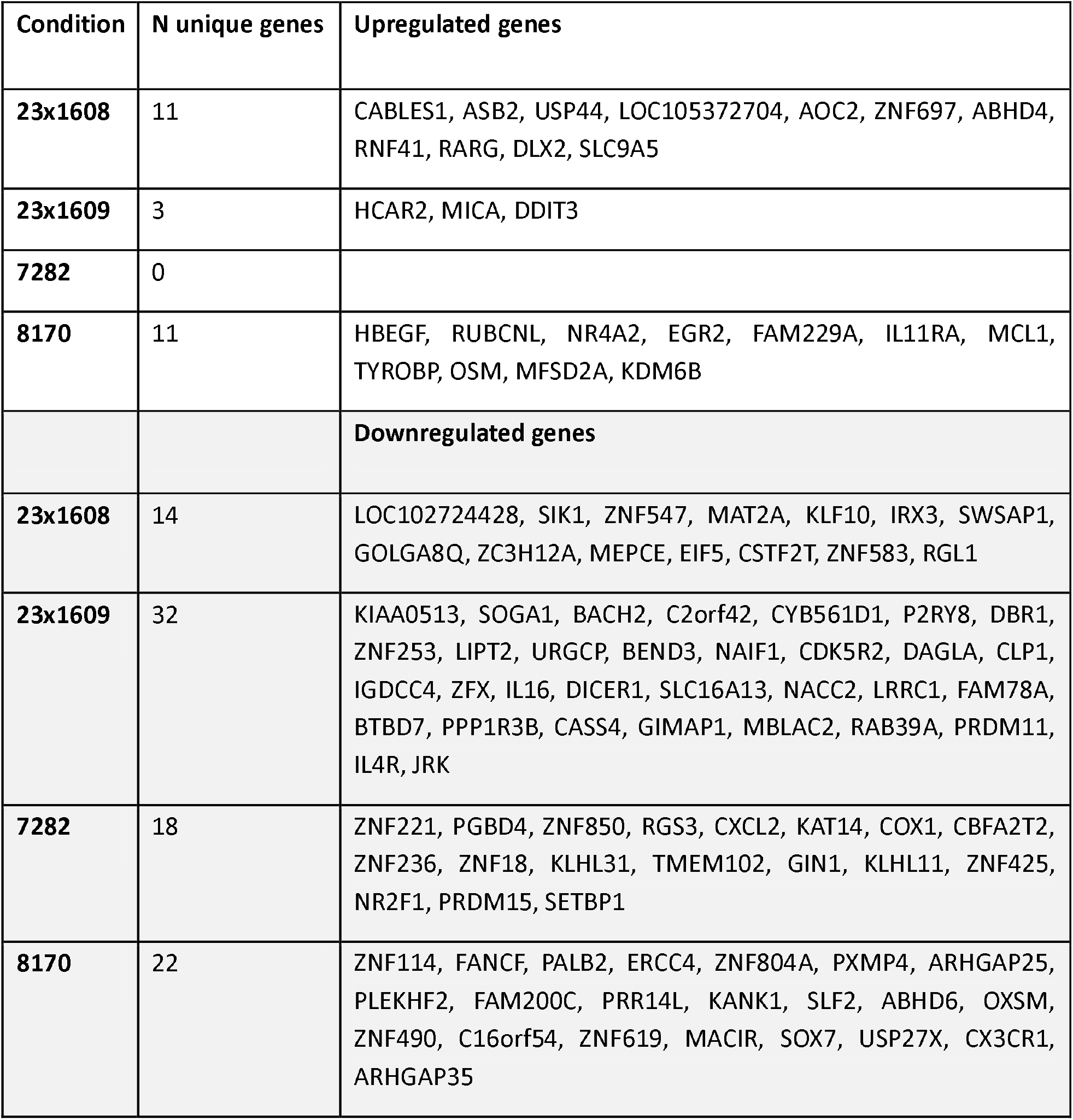
Unique genes up- or downregulated in U937 monocytes in response to each isolate at 6 hpi, MOI 50, after washing at 1 hpi.

**Figure 2:**
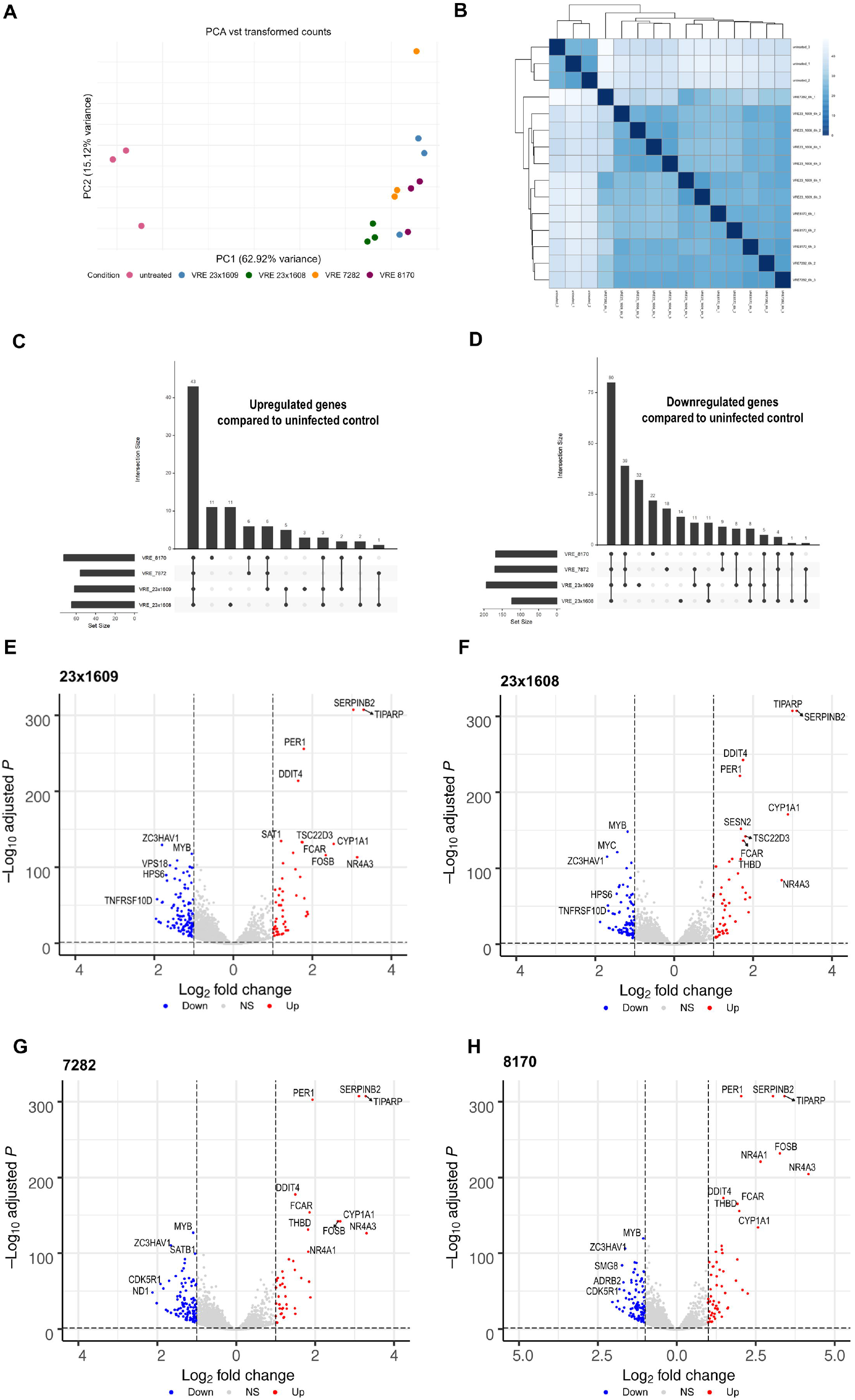
Differential gene regulation of U937 cells in response to *VREfm* at 6 hpi. U937 cells were infected with the *VREfm* isolates 23×1609, 23×1608, 7282, and 8170 at MOI 50 and washed at 2 hpi. RNA was isolated at 6 hpi. **(A)** Principal component analysis based on variance-stabilizing transformed (vst) counts using the top 1000 most variable protein-coding genes of uninfected U937 cells and those infected with the different *VREfm* isolates. **(B)** Sample (Euclidean) distance heatmap of each sample replicate based on DESeq2 protein-coding vst counts. UpSet plots depict overlap of significantly, at least two-fold **(C)** upregulated and **(D)** downregulated genes across conditions. **(E-H)** Volcano plots of cells infected with *VREfm* isolates compared to uninfected control cells delineate homogenous differential transcriptional regulation (padj < 0.05, | Log2FC | > 1).

Network analysis of the shared core gene sets (**Fig. 3A**) emphasized a key role of TNF in linking various downregulated gene networks. A majority of pathways are shared across isolates, as depicted by the overlap of upregulated versus downregulated genes (purple lines) and pathways (blue lines) according to GO:BP terminology across isolates (**Fig. 3B**). We show differential activation of various pathways involved in inflammation, cellular differentiation, and regulation of programmed cell death, likely affecting cellular homeostasis upon persistent (intracellular) infection.

**Figure 3:**
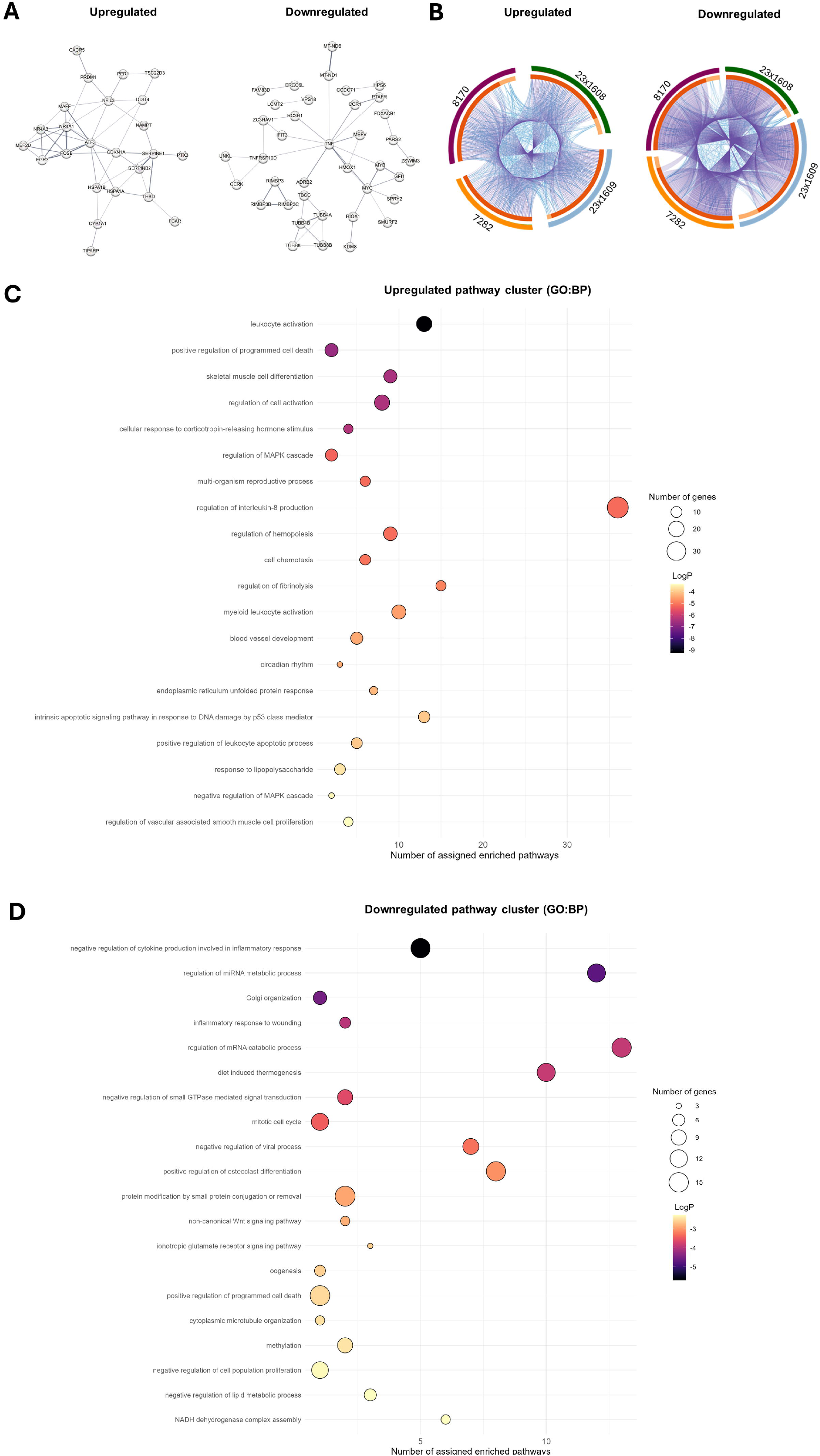
Functional enrichment of DEGs in U937 cells infected with *VREfm* at 6 hpi. U937 cells were infected with the *VREfm* isolates 23×1608, 23×1609, 7282, and 8170 at MOI 50 and washed at 2 hpi. RNA was isolated at 6 hpi. **(A)** Full STRING network of up- and downregulated protein-coding genes (padj < 0.05, | Log2FC | > 1) in U937 cells shared across all four *VREfm* isolates at 6 hpi (edges indicate confidence; medium confidence 0.400), disconnected nodes not shown. (B) Circos plots indicate overlap of gene lists only at the gene (purple curves) and pathway (blue curves; GO:BP term) level. Significantly (padj < 0.05), at least two-fold (C) upregulated or (D) downregulated protein-coding genes were analyzed for enriched pathway clusters (GO:BP) using Metascape for multiple gene lists (p < 0.01; minimal overlap = 3, enrichment factor = 1.5), showing the driver pathway’s adjusted enrichment score (LogP), gene set size (bubble size) and number of pathways assigned to each cluster (x axis).

The overall top 20 upregulated (**Fig. 3C**) and downregulated (**Fig. 3D**) pathway clusters across isolates revealed e.g. proinflammatory, cell proliferation, and compartmentalization pathways to be highly regulated. Relative expression of specific genes involved in each cluster showed some variation between the different *VREfm* isolates (**Fig. 4**). The top five regulated clusters were (up): “*leukocyte activation*”, “*positive regulation of programmed cell death*” and “*skeletal muscle cell differentiation*”, “*regulation of cell activation*” and “*cellular response to corticotropin-releasing hormone stimulus*”; (down): “*negative regulation of cytokine production involved in inflammatory response*”, “*regulation of miRNA metabolic process*”, “*Golgi organization*”, “*inflammatory response to wounding*” and “*regulation of mRNA catabolic process*”.

**Figure 4:**
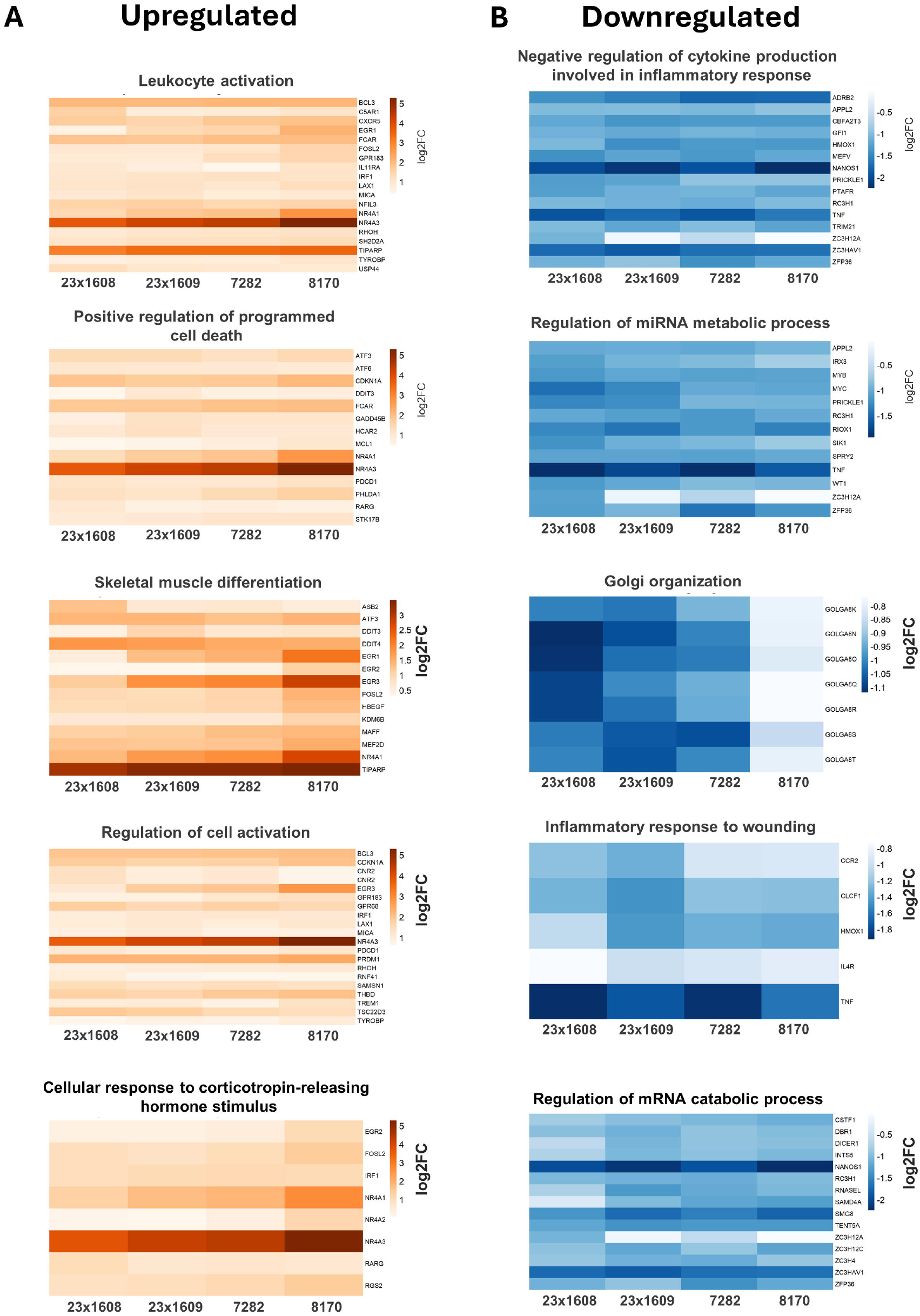
Top 5 enriched pathway clusters and relative expression of included DEGs for **(A)** upregulated and **(B)** downregulated clusters in uninfected U937 cells or those infected with each *VREfm* isolate at 6 hpi. Significantly (padj ≤ 0.05), at least two-fold differentially regulated, protein-coding genes were included and enrichment was performed using Metascape for multiple gene lists (p < 0.01; minimal overlap = 3, enrichment factor = 1.5).

**Figure S2:**
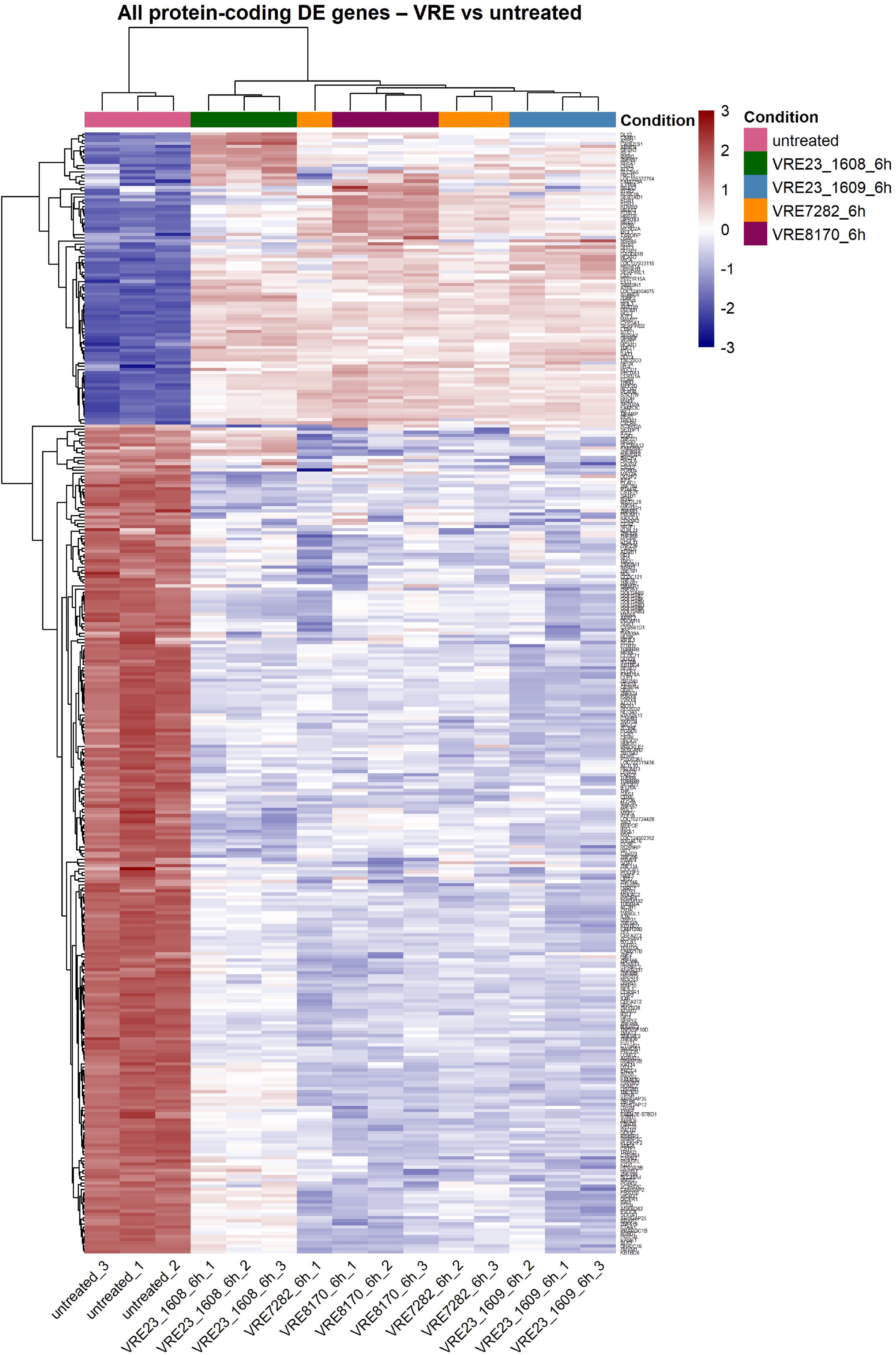
Z-scores of all protein-coding DEGs. (padj ≤0.05; | Log2FC | ≥1) in uninfected U937 cells or those infected with different *VREfm* isolates in triplicate at 6 hpi. Euclidean distances and Z-scores are based on DESeq2 protein-coding vst-counts for each sample replicate.

**Figure S3:**
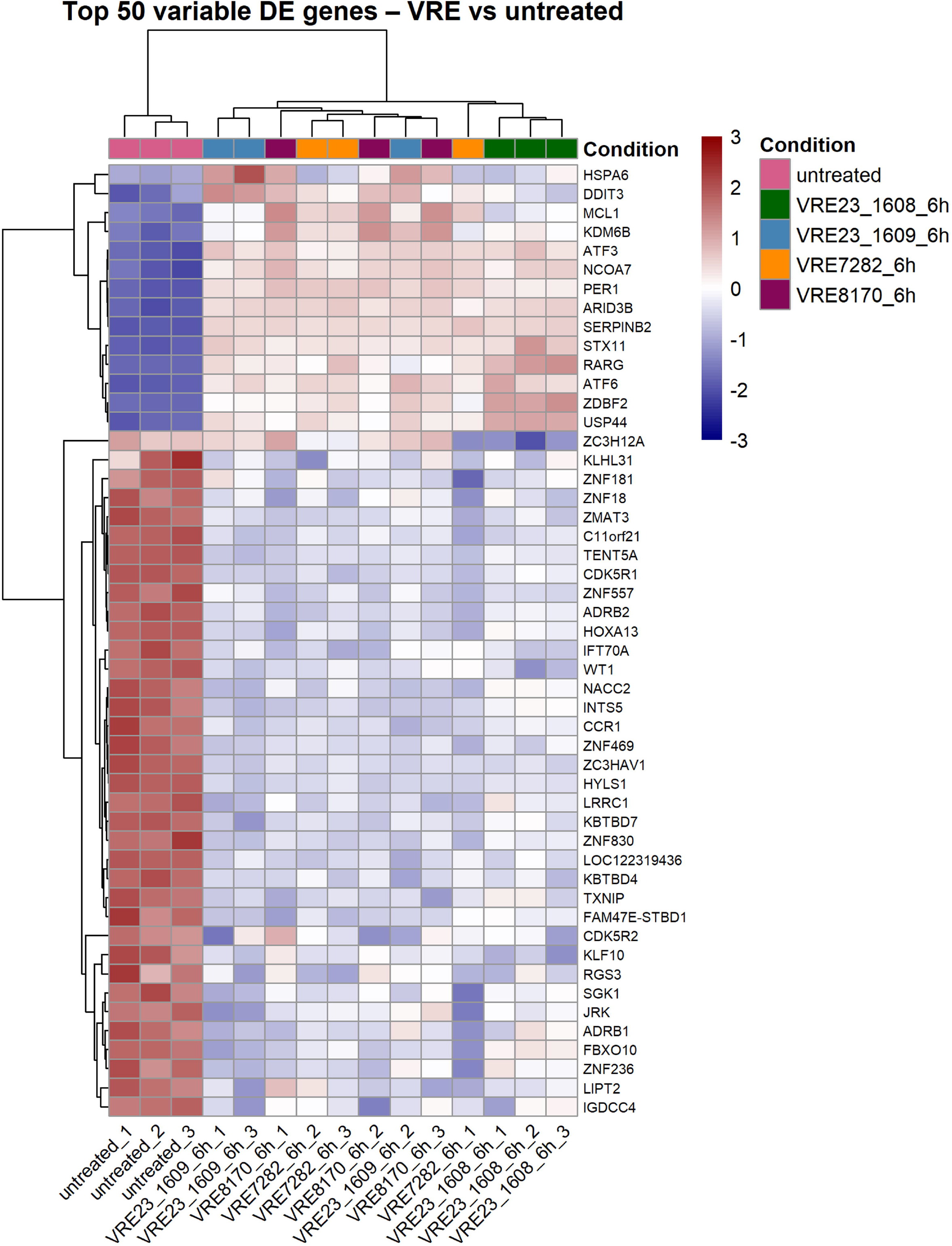
Z-scores of selected top 50 DEGs showing the strongest variation across samples in uninfected U937 cells or those infected with different *VREfm* isolates in triplicate at 6 hpi. Significant (padj ≤0.05), at least two-fold differentially regulated, protein-coding genes were included. Euclidean distances and Z-scores are based on DESeq2 protein-coding vst-counts for each sample replicate.

In contrast to *S. aureus*-infected U937 cells (Walter et al., 2026), the host’s transcriptional alterations were homogenous across the different *VREfm* isolates despite certain genomic variability. The composition of the observed host response suggests a model in which *VREfm* infection drives the host cells into a robust pro-inflammatory and stress-adapted state characterized by innate immune activation, rapid transcriptional reprogramming, and partial relief of post-transcriptional inhibitory control. Simultaneously, the concomitant downregulation of inflammatory signatures, e.g. TNF and related pathways, suggests a dynamic cross-talk between pro-inflammatory activation and host defense feedback regulation, which is typical of host-pathogen interactions in myeloid cells (Das et al., 2015; Vannella and Wynn, 2017). This transcriptional balance may reflect both the immediate immune response to microbial challenge and compensatory mechanisms that limit excessive inflammatory damage. Especially in nosocomial *Efm* isolates, Toll/interleukin-1 receptor-(TIR)-domain containing proteins have been shown to be expressed during BSIs and to impede immune cell activation e.g. by regulating NF-κB activity (Zou et al., 2014). These secreted proteins do not necessarily require cell-cell contact and have been described in other species (Askarian et al., 2014; Rana et al., 2013; Wagner et al., 2018). As we merely investigated the host response, dual RNA sequencing will reveal the interplay of virulence factors and transcriptional regulation. Additionally, responses of the U937 cell line may not accurately translate to primary cells and *in vivo* conditions, which remains a major limitation of this study.

Another major limitation is based on available experimental designs allowing for the investigation of host effects induced by intracellular bacteria only, e.g. gentamicin protection assays, due to isolate-dependent antimicrobial resistances to gentamicin, linezolid, combined linezolid and gentamicin, or daptomycin at concentrations that were still tolerable to host cells (not shown). Consequently, based on the washing step at 1 hpi, this set-up aimed to concentrate rather than limit the observed effects on internalized bacteria or those with direct host cell contact. However, we cannot exclude a non-negligible impact of extracellular bacteria that remained after washing or were released throughout the course of infection. To further investigate *VREfm* as an intracellular pathogen, alternatives to the gentamicin protection assay used in our study should be implemented, as high antibiotic concentrations may also affect their survival within the host cell. This may include the use of rapid-acting agents such as antimicrobial peptides or phage-derived endolysins, especially when working with otherwise antibiotic-resistant bacteria. Additionally, other pathogens such as *Efc* should be used as controls.

Taken together, our study underscores the need for more detailed analyses of *VREfm – host* crosstalk, with a particular focus on whether enterococci establish intracellular survival niches in different human cell types and tissues, potentially promoting their capacity to cause invasive, systemic infections. We describe close association of several *VREfm* isolates with human U937 monocytes and features compatible with their internalization. Under the specific conditions employed in this study, host cell death was induced only at later infection stages at isolate-dependent levels, but was abrogated by bc insertion preventing direct host cell contact or internalization. Conversely, TNF-R1 receptor deficiency strongly promoted host cell cytotoxicity. Transcriptional reprogramming of U937 cells, widely consistent across the different *VREfm* isolates, resulted in early monocyte activation characterized by innate sensing, immediate early transcription, and stress adaptation pathways. Pathways that usually constrain cytokine abundance and RNA stability were suppressed.

Future work will disentangle the relationship between host cell activation, internalization and direct host-pathogen contact and TNF/TNF-R1 controlled host cell fates.

## Authors’ contributions

Experiments: AW, TP, IG, JF. Data analysis: AW, JF, TB, TG. Conceptualization: AW, JF. Manuscript draft: AW, JF. Manuscript editing: AW, TP, TB, IG, TG, BK, WSB, AR, JF.

## Data and materials availability

All data needed to evaluate the conclusions in this paper are present either in the main text or in the supplementary materials. RNA-seq data have been submitted together with data from an *S. aureus-centric* study (Walter et al., 2026) and are available at https://www.ncbi.nlm.nih.gov/geo/query/acc.cgi?acc=GSE328149.

## Acknowledgment

Library preparation and mRNAseq were performed at the Genomics Core Facility “KFB - Center of Excellence for Fluorescent Bioanalytics” (University of Regensburg, Regensburg, Germany; www.kfb-regensburg.de).

## References

Archambaud, C., Nunez, N., da Silva, R.A.G., Kline, K.A., and Serror, P. (2024). Enterococcus faecalis: an overlooked cell invader. Microbiol Mol Biol Rev 88, e0006924, 10.1128/mmbr.00069-24.

Askarian, F., van Sorge, N.M., Sangvik, M., Beasley, F.C., Henriksen, J.R., Sollid, J.U., van Strijp, J.A., Nizet, V., and Johannessen, M. (2014). A Staphylococcus aureus TIR domain protein virulence factor blocks TLR2-mediated NF-kappaB signaling. Journal of innate immunity 6, 485–498, 10.1159/000357618.

Ayobami, O., Willrich, N., Reuss, A., Eckmanns, T., and Markwart, R. (2020). The ongoing challenge of vancomycin-resistant Enterococcus faecium and Enterococcus faecalis in Europe: an epidemiological analysis of bloodstream infections. Emerg Microbes Infect 9, 1180–1193, 10.1080/22221751.2020.1769500.

Baldassarri, L., Bertuccini, L., Creti, R., Filippini, P., Ammendolia, M.G., Koch, S., Huebner, J., and Orefici, G. (2005). Glycosaminoglycans mediate invasion and survival of Enterococcus faecalis into macrophages. J Infect Dis 191, 1253–1262, 10.1086/428778.

Benes, J., Dzupova, O., Setina, M., Feuereisl, R., Svec, P., and Pantucek, R. (2013). Relapsing endocarditis caused by Enterococcus faecalis forming small colony variants. Scand J Infect Dis 45, 800–803, 10.3109/00365548.2013.800227.

Brinkwirth, S., Feig, M., Noll, I., Eckmanns, T., Dorre, A., Haller, S., and Willrich, N. (2025). Changing dynamics of bloodstream infections due to methicillin-resistant Staphylococcus aureus and vancomycin-resistant Enterococcus faecium in Germany, 2017-2023: a continued burden of disease approach. Antimicrob Resist Infect Control 14, 4, 10.1186/s13756-025-01522-9.

Cai, J., Zhou, H., Liu, M., Zhang, D., Lv, J., Xue, H., Zhou, H., and Zhang, W. (2025). Host immunity and intracellular bacteria evasion mechanisms: Enhancing host-directed therapies with drug delivery systems. Int J Antimicrob Agents 65, 107492, 10.1016/j.ijantimicag.2025.107492.

Cairns, K.A., Udy, A.A., Peel, T.N., Abbott, I.J., Dooley, M.J., and Peleg, A.Y. (2023). Therapeutics for Vancomycin-Resistant Enterococcal Bloodstream Infections. Clin Microbiol Rev 36, e0005922, 10.1128/cmr.00059-22.

Canadian Nosocomial Infection Surveillance, P. (2026). Healthcare-associated infections and antimicrobial resistance in Canadian acute care hospitals, 2020-2024. Can Commun Dis Rep 52, 205–222, 10.14745/ccdr.v52i05a05.

Caplunik-Pratsch, A., Kieninger, B., Donauer, V.A., Brauer, J.M., Meier, V.M.K., Seisenberger, C., Rath, A., Loibl, D., Eichner, A., Fritsch, J., and Schneider-Brachert, W. (2024). Introduction and spread of vancomycin-resistant Enterococcus faecium (VREfm) at a German tertiary care medical center from 2004 until 2010: a retrospective whole-genome sequencing (WGS) study of the molecular epidemiology of VREfm. Antimicrob Resist Infect Control 13, 20, 10.1186/s13756-024-01379-4.

Caplunik-Pratsch, A., Kieninger, B., Hansch, S., Rath, A., Fritsch, J., Eichner, A., Hanses, F., Schneider-Brachert, W., and Hitzenbichler, F. (2025). The potential of genome-oriented blood culture surveillance of vancomycin-resistant enterococcus faecium (VRE) to mirror local VRE epidemiology: a retrospective analysis and systematic comparison of VRE blood culture and VRE first patient isolates. BMC Infect Dis 25, 1685, 10.1186/s12879-025-12183-9.

Choi, E.J., Iwasa, M., Han, K.I., Kim, W.J., Tang, Y., Hwang, Y.J., Chae, J.R., Han, W.C., Shin, Y.S., and Kim, E.K. (2016). Heat-Killed Enterococcus faecalis EF-2001 Ameliorates Atopic Dermatitis in a Murine Model. Nutrients 8, 146, 10.3390/nu8030146.

da Silva, R.A.G., Tay, W.H., Ho, F.K., Tanoto, F.R., Chong, K.K.L., Choo, P.Y., Ludwig, A., and Kline, K.A. (2022). Enterococcus faecalis alters endo-lysosomal trafficking to replicate and persist within mammalian cells. PLoS pathogens 18, e1010434, 10.1371/journal.ppat.1010434.

Dai, X., Deng, Z., Liang, Y., Chen, L., Jiang, W., and Zhao, W. (2020). Enterococcus faecalis induces necroptosis in human osteoblastic MG63 cells through the RIPK3/MLKL signalling pathway. Int Endod J 53, 1204–1215, 10.1111/iej.13323.

Das, A., Sinha, M., Datta, S., Abas, M., Chaffee, S., Sen, C.K., and Roy, S. (2015). Monocyte and macrophage plasticity in tissue repair and regeneration. The American journal of pathology 185, 2596–2606, 10.1016/j.ajpath.2015.06.001.

Deneke, C. (2020). MiLongA: A snakemake workflow for long-read assembly. gitlab.

Di Giacinto, C., Marinaro, M., Sanchez, M., Strober, W., and Boirivant, M. (2005). Probiotics ameliorate recurrent Th1-mediated murine colitis by inducing IL-10 and IL-10-dependent TGF-beta-bearing regulatory cells. Journal of immunology 174, 3237–3246, 10.4049/jimmunol.174.6.3237.

Dreschers, S., Gille, C., Haas, M., Seubert, F., Platen, C., and Orlikowsky, T.W. (2017). Reduced internalization of TNF-a/TNFR1 down-regulates caspase dependent phagocytosis induced cell death (PICD) in neonatal monocytes. PloS one 12, e0182415, 10.1371/journal.pone.0182415.

Edelmann, B., Bertsch, U., Tchikov, V., Winoto-Morbach, S., Perrotta, C., Jakob, M., Adam-Klages, S., Kabelitz, D., and Schütze, S. (2011). Caspase-8 and caspase-7 sequentially mediate proteolytic activation of acid sphingomyelinase in TNF-R1 receptosomes. The EMBO journal 30, 379–394, 10.1038/emboj.2010.326.

Egido, S.H., Ruiz, M.S., Ines Revuelta, S., Garcia, I.G., and Bellido, J.L. (2016). Enterococcus faecium small colony variant endocarditis in an immunocompetent patient. New Microbes New Infect 9, 47–49, 10.1016/j.nmni.2015.11.006.

Fiore, E., Van Tyne, D., and Gilmore, M.S. (2019). Pathogenicity of Enterococci. Microbiol Spectr 7, 10.1128/microbiolspec.GPP3-0053-2018.

Fraunholz, M., and Sinha, B. (2012). Intracellular Staphylococcus aureus: live-in and let die. Front Cell Infect Microbiol 2, 43, 10.3389/fcimb.2012.00043.

Fritsch, J., Fickers, R., Klawitter, J., Sarchen, V., Zingler, P., Adam, D., Janssen, O., Krause, E., and Schütze, S. (2016). TNF induced cleavage of HSP90 by cathepsin D potentiates apoptotic cell death. Oncotarget 7, 75774–75789, 10.18632/oncotarget.12411.

Fritsch, J., Stephan, M., Tchikov, V., Winoto-Morbach, S., Gubkina, S., Kabelitz, D., and Schutze, S. (2014). Cell fate decisions regulated by K63 ubiquitination of tumor necrosis factor receptor 1. Molecular and cellular biology 34, 3214–3228, 10.1128/MCB.00048-14.

Fu, Q., Yuan, J., Wang, L., Ran, H., Li, F., Liu, F., Zhang, J., Liu, W., Huang, W., Huang, Y., and Xia, X. (2020). Proteomic analysis of murine macrophages mitochondria and lysosomes reveal Cathepsin D as a potential broad-spectrum antimicrobial protein. Journal of proteomics 223, 103821, 10.1016/j.jprot.2020.103821.

Gröbner, S., Beck, J., Schaller, M., Autenrieth, I.B., and Schulte, B. (2012). Characterization of an Enterococcus faecium small-colony variant isolated from blood culture. Int J Med Microbiol 302, 40–44, 10.1016/j.ijmm.2011.07.001.

Heinrich, M., Neumeyer, J., Jakob, M., Hallas, C., Tchikov, V., Winoto-Morbach, S., Wickel, M., Schneider-Brachert, W., Trauzold, A., Hethke, A., and Schütze, S. (2004). Cathepsin D links TNF-induced acid sphingomyelinase to Bid-mediated caspase-9 and -3 activation. Cell death and differentiation 11, 550–563, 10.1038/sj.cdd.4401382.

Horing, S., Schutz, M., Autenrieth, I.B., and Grobner, S. (2012). Lysozyme facilitates adherence of Enterococcus faecium to host cells and induction of necrotic cell death. Microbes and infection/Institut Pasteur 14, 554–562, 10.1016/j.micinf.2012.01.005.

Höring, S., Sobotta, K., Schneider, S., Löffler, B., and Rodel, J. (2019). Dwarfs in disguise: multiple spinal abscesses and spondylodiscitis caused by an Enterococcus faecium small-colony variant. Access Microbiol 1, e000012, 10.1099/acmi.0.000012.

Horsley, H., Malone-Lee, J., Holland, D., Tuz, M., Hibbert, A., Kelsey, M., Kupelian, A., and Rohn, J.L. (2013). Enterococcus faecalis subverts and invades the host urothelium in patients with chronic urinary tract infection. PloS one 8, e83637, 10.1371/journal.pone.0083637.

Kahl, B.C., Becker, K., and Loffler, B. (2016). Clinical Significance and Pathogenesis of Staphylococcal Small Colony Variants in Persistent Infections. Clin Microbiol Rev 29, 401–427, 10.1128/CMR.00069-15.

Kao, P.H.N., and Kline, K.A. (2019). Dr. Jekyll and Mr. Hide: How Enterococcus faecalis Subverts the Host Immune Response to Cause Infection. Journal of molecular biology 431, 2932–2945, 10.1016/j.jmb.2019.05.030.

Leding, C., Meddis, A., Holler, J.G., Burisch, J., Kragstrup, T.W., Skov, L., and Benfield, T. (2026). Tumour necrosis factor-alpha inhibitors, immune-mediated inflammatory diseases, and bloodstream infections: results from a nationwide case-control study. Clin Microbiol Infect 32, 951–958, 10.1016/j.cmi.2026.02.005.

Li, X., Korner, H., and Liu, X. (2020). Susceptibility to Intracellular Infections: Contributions of TNF to Immune Defense. Front Microbiol 11, 1643, 10.3389/fmicb.2020.01643.

Li, Y., Tong, Z., and Ling, J. (2019). Effect of the three Enterococcus faecalis strains on apoptosis in MC3T3 cells. Oral Dis 25, 309–318, 10.1111/odi.12883.

Millan, D., Chiriboga, C., Patarroyo, M.A., and Fontanilla, M.R. (2013). Enterococcus faecalis internalization in human umbilical vein endothelial cells (HUVEC). Microb Pathog 57, 62–69, 10.1016/j.micpath.2012.11.007.

Morgan, M.J., and Liu, Z.G. (2010). Reactive oxygen species in TNFalpha-induced signaling and cell death. Mol Cells 30, 1–12, 10.1007/s10059-010-0105-0.

Ogihara, S., Saito, R., Sawabe, E., Hagihara, M., and Tohda, S. (2016). First Japanese case of infectious endocarditis due to Enterococcus faecalis small-colony variants. J Infect Chemother 22, 716–719, 10.1016/j.jiac.2016.03.010.

Rana, R.R., Zhang, M., Spear, A.M., Atkins, H.S., and Byrne, B. (2013). Bacterial TIR-containing proteins and host innate immune system evasion. Med Microbiol Immunol 202, 1–10, 10.1007/s00430-012-0253-2.

Rath, A., Kieninger B, Mirzaliyeva N, Werner G, Bender JK, Fische MA, Cabal-Rosel A, Ruppitsch W, Seth-Smith HMB, Egli A, Halabi M, Hörtenhuber A, Prammer W, Salaheddin Y, Kerschner H, Hartl R, Ehrenschwender M, Ambrosch A, Kalinowski J, Klages LJ, Rückert-Reed C, Busche T, Kratzer A, Caplunik-Pratsch A, Eichner A, Fritsch J, and W., S.-B. (2025). Spread of the novel vancomycin-resistant Enterococcus faecium strain ST1299/vanA from local level in Germany to cross-border level in Austria, 2018 to 2022. Euro Surveill 30, 10.2807/1560-7917.ES.2025.30.20.2400389.

Rath, A., and Schneider-Brachert, W. (2026). Limited, outbreak-biased and unevenly reported: genomic insights into vancomycin-resistant Enterococcus faecium in Germany as reported between 2015 and 2025. Infection, 10.1007/s15010-026-02890-z.

Schneider-Brachert, W., Tchikov, V., Merkel, O., Jakob, M., Hallas, C., Kruse, M.L., Groitl, P., Lehn, A., Hildt, E., Held-Feindt, J., Dobner, T., Kabelitz, D., Krönke, M., and Schütze, S. (2006). Inhibition of TNF receptor 1 internalization by adenovirus 14.7K as a novel immune escape mechanism. The Journal of clinical investigation 116, 2901–2913, 10.1172/JCI23771.

Schramm, M., Herz, J., Haas, A., Kronke, M., and Utermohlen, O. (2008). Acid sphingomyelinase is required for efficient phago-lysosomal fusion. Cellular microbiology 10, 1839–1853, 10.1111/j.1462-5822.2008.01169.x.

Sussmuth, S.D., Muscholl-Silberhorn, A., Wirth, R., Susa, M., Marre, R., and Rozdzinski, E. (2000). Aggregation substance promotes adherence, phagocytosis, and intracellular survival of Enterococcus faecalis within human macrophages and suppresses respiratory burst. Infect Immun 68, 4900–4906, 10.1128/IAI.68.9.4900-4906.2000.

Szperlinski, M., Schermuly, E., Luhrmann, A., and Kozjak-Pavlovic, V. (2025). Obligate intracellular bacteria and host cell death pathways-the matter of life and death. Cell Tissue Res 402, 413–439, 10.1007/s00441-025-04014-9.

Tatusova, T., DiCuccio, M., Badretdin, A., Chetvernin, V., Nawrocki, E.P., Zaslavsky, L., Lomsadze, A., Pruitt, K.D., Borodovsky, M., and Ostell, J. (2016). NCBI prokaryotic genome annotation pipeline. Nucleic acids research 44, 6614–6624, 10.1093/nar/gkw569.

Thakur, A., Mikkelsen, H., and Jungersen, G. (2019). Intracellular Pathogens: Host Immunity and Microbial Persistence Strategies. J Immunol Res 2019, 1356540, 10.1155/2019/1356540.

Thieme, L., Mirza, K.A., Azimi, M., Enslinger, N., Lohde, M., Jundzill, M., Brandt, C., Pletz, M.W., and Makarewicz, O. (2026). Extended lag phases and variable antibiotic tolerance in beta-lactam-induced small colony variants of Enterococcus faecalis. BMC Microbiol 26, 10.1186/s12866-026-05403-y.

Vannella, K.M., and Wynn, T.A. (2017). Mechanisms of Organ Injury and Repair by Macrophages. Annu Rev Physiol 79, 593–617, 10.1146/annurev-physiol-022516-034356.

Wagner, T.M., Janice, J., Paganelli, F.L., Willems, R.J., Askarian, F., Pedersen, T., Top, J., de Haas, C., van Strijp, J.A., Johannessen, M., and Hegstad, K. (2018). Enterococcus faecium TIR-Domain Genes Are Part of a Gene Cluster Which Promotes Bacterial Survival in Blood. Int J Microbiol 2018, 1435820, 10.1155/2018/1435820.

Waguia Kontchou, C., Tzivelekidis, T., Gentle, I.E., and Hacker, G. (2016). Infection of epithelial cells with Chlamydia trachomatis inhibits TNF-induced apoptosis at the level of receptor internalization while leaving non-apoptotic TNF-signalling intact. Cellular microbiology 18, 1583–1595, 10.1111/cmi.12598.

Walter, A., Bischler, T., Jungblut, M., Breitsprecher, L.F., Vilotic, F., Beck, J., Hofmann, L., Schafer, N., Ziesmann, T., Franzkoch, R., Harteis, S., Gadjalova, I., Distler, U., Beliu, G., Psathaki, O.E., Hensel, M., Schneider-Brachert, W., Grafenhan, T., Stempfl, T., Kieninger, B., Muhlen, S., Alt, V., Mannala, G.K., and Fritsch, J. (2026). Intracellular Staphylococcus aureus triggers isolate-specific host transcriptional responses alongside TNF-R1 regulated cell fate decisions. Cell communication and signaling : CCS 24, 10.1186/s12964-026-03246-8.

Wei, Y., Palacios Araya, D., and Palmer, K.L. (2024). Enterococcus faecium: evolution, adaptation, pathogenesis and emerging therapeutics. Nat Rev Microbiol 22, 705–721, 10.1038/s41579-024-01058-6.

Wellinghausen, N., Chatterjee, I., Berger, A., Niederfuehr, A., Proctor, R.A., and Kahl, B.C. (2009). Characterization of clinical Enterococcus faecalis small-colony variants. J Clin Microbiol 47, 2802–2811, 10.1128/JCM.00485-09.

Wells, C.L., Moore, E.A., Hoag, J.A., Hirt, H., Dunny, G.M., and Erlandsen, S.L. (2000). Inducible expression of Enterococcus faecalis aggregation substance surface protein facilitates bacterial internalization by cultured enterocytes. Infect Immun 68, 7190–7194, 10.1128/IAI.68.12.7190-7194.2000.

WHO (2024). WHO Bacterial Priority Pathogens List, 2024.

Xiong, X., Tian, S., Yang, P., Lebreton, F., Bao, H., Sheng, K., Yin, L., Chen, P., Zhang, J., Qi, W., Ruan, J., Wu, H., Chen, H., Breault, D.T., Wu, H., Earl, A.M., Gilmore, M.S., Abraham, J., and Dong, M. (2022). Emerging enterococcus pore-forming toxins with MHC/HLA-I as receptors. Cell 185, 1157–1171 e1122, 10.1016/j.cell.2022.02.002.

Yazdanpanah, B., Wiegmann, K., Tchikov, V., Krut, O., Pongratz, C., Schramm, M., Kleinridders, A., Wunderlich, T., Kashkar, H., Utermohlen, O., Bruning, J.C., Schutze, S., and Kronke, M. (2009). Riboflavin kinase couples TNF receptor 1 to NADPH oxidase. Nature 460, 1159–1163, 10.1038/nature08206.

Zhang, S., Lebreton, F., Mansfield, M.J., Miyashita, S.I., Zhang, J., Schwartzman, J.A., Tao, L., Masuyer, G., Martinez-Carranza, M., Stenmark, P., Gilmore, M.S., Doxey, A.C., and Dong, M. (2018). Identification of a Botulinum Neurotoxin-like Toxin in a Commensal Strain of Enterococcus faecium. Cell host & microbe 23, 169–176 e166, 10.1016/j.chom.2017.12.018.

Zhou, Y., Zhou, B., Pache, L., Chang, M., Khodabakhshi, A.H., Tanaseichuk, O., Benner, C., and Chanda, S.K. (2019). Metascape provides a biologist-oriented resource for the analysis of systems-level datasets. Nature communications 10, 1523, 10.1038/s41467-019-09234-6.

Zingler, P., Särchen, V., Glatter, T., Caning, L., Saggau, C., Kathayat, R.S., Dickinson, B.C., Adam, D., Schneider-Brachert, W., Schütze, S., and Fritsch, J. (2019). Palmitoylation is required for TNF-R1 signaling. Cell communication and signaling : CCS 17, 90, 10.1186/s12964-019-0405-8.

Zou, J., Baghdayan, A.S., Payne, S.J., and Shankar, N. (2014). A TIR domain protein from E. faecalis attenuates MyD88-mediated signaling and NF-kappaB activation. PloS one 9, e112010, 10.1371/journal.pone.0112010.

Zou, J., and Shankar, N. (2014). Enterococcus faecalis infection activates phosphatidylinositol 3-kinase signaling to block apoptotic cell death in macrophages. Infect Immun 82, 5132–5142, 10.1128/IAI.02426-14.

